# PixelDeck: A local-first media library manager for biomedical imaging

**DOI:** 10.64898/2026.04.24.719925

**Authors:** Benjamin L. Kidder

**Author notes:** Correspondence: Benjamin L. Kidder.

## Abstract

Modern biomedical imaging workflows generate large volumes of derived images and short videos that must be reviewed, compared, curated, and reused following primary acquisition and analysis. In practice, these assets are often dispersed across nested filesystem hierarchies on local drives, external media, or network storage, limiting efficient retrieval, deduplication, and figure assembly. We present PixelDeck, an open-source, local-first browser application for organizing and interactively browsing large biomedical image and video libraries on commodity workstations. PixelDeck integrates recursive folder import, SHA-256-based duplicate detection, metadata extraction, thumbnail and preview generation, full-text search, and asynchronous export within a responsive interface, supported by a modular ingestion pipeline, managed storage layer, and interactive browsing environment optimized for high-volume media collections. The system is implemented using a Next.js and React frontend, a SQLite metadata store accessed via Prisma, managed local media storage, and a background worker that executes import and export tasks asynchronously, enabling scalable processing on standard hardware. To evaluate performance, we conducted structured benchmark imports using public histopathology images curated from PanopTILs, SICAPv2, and PanNuke datasets, where dataset-specific import behavior, duplicate detection, and ingestion metrics were recorded as reproducible outputs. Embedding-based analysis further demonstrates dataset-level separation consistent with underlying image characteristics. These results show that PixelDeck provides an efficient, scalable local curation layer for heterogeneous biomedical imaging collections, enabling streamlined dataset exploration and preparation for downstream analysis.

## INTRODUCTION

Biomedical imaging has become increasingly data-intensive, with modern workflows producing large numbers of processed image derivatives, multichannel composites, segmentation overlays, quality-control snapshots, time-series exports, and assay review videos^1-7^. While acquisition and analytic pipelines are often specialized and well developed, downstream management of these derived outputs is frequently handled through conventional filesystem browsing. In practice, this creates a persistent gap between image generation and image reuse, particularly when investigators need to retrieve prior outputs, identify duplicates, compare results across experiments, or assemble reproducible subsets for downstream analysis and manuscript preparation^1-4, 8-11^.

Existing imaging platforms address important parts of this broader ecosystem, including multidimensional image modeling, image archive management, and integrated bioimage analysis environments^4, 8-11^. However, many laboratories still require a lighter-weight tool for local curation of exported media collections rather than full acquisition management or large-scale server deployment. This need is especially relevant for review-oriented datasets derived from histology, fluorescence microscopy, and high-throughput imaging workflows, where large collections of images or short video outputs accumulate across nested directory structures on local and removable storage. These data are valuable for interpretation and communication but often difficult to search, deduplicate, and repurpose efficiently once they leave the original acquisition or analysis context^1-3, 5, 6^.

PixelDeck was developed to address this practical gap. The software is intended as a local-first curation and retrieval layer for large biomedical media libraries rather than a replacement for instrument-native image-management systems or laboratory information management systems. Its design emphasizes transparent file handling, reproducible metadata storage, responsive browser-based interaction, and minimal infrastructure requirements. By coupling content-based deduplication, generated derivative media, full-text indexing, and asynchronous processing on commodity workstations, PixelDeck provides a pragmatic approach for managing exported image and video assets in biomedical research workflows^12^.

Here we describe the architecture, implementation, and intended use of PixelDeck, and discuss its current scope and limitations. In addition, we incorporate a public-dataset evaluation framework based on benchmark imports from histopathology subcollections. To further characterize dataset structure, we apply dimensionality reduction using PCA and UMAP to imported image features, enabling visualization of dataset-level organization and separation as part of the evaluation framework.

## METHODS

PixelDeck follows a modular local-first design composed of a browser-facing application layer, a structured metadata layer, a managed filesystem storage layer, and a dedicated background worker for asynchronous processing. The frontend is implemented with Next.js and React and provides the main user interface for grid and table browsing, search, filtering, selection, collection management, and export initiation. HTTP API routes exposed by the application coordinate interactions among the user interface, the metadata store, and the worker-driven job system.

Metadata persistence is implemented with SQLite and accessed through Prisma. The schema is centered on an Asset model that stores descriptive and technical information including original filename, display name, folder context, source label, media type, dimensions, duration, file size, SHA-256 hash, timestamps, and paths to original and derived media. Additional tables support tags, collections, import jobs, duplicate events, export jobs, saved searches, and application settings. SQLite full-text search is used to support retrieval across filenames, display names, sources, tags, and aggregated searchable text. This approach combines a lightweight relational database with local query flexibility and transparent file-level storage^4, 8-11^.

For benchmark evaluation and downstream feature-space analysis, we focused on RGB image subsets from PanopTILs, SICAPv2, and PanNuke^13-15^. Each source folder was imported independently under a distinct PixelDeck source label so that discovered files, processed assets, duplicate events, skipped files, failed files, and job status could be tracked separately.

Imported media are normalized into a managed directory structure containing immutable originals and separate directories for thumbnails, previews, video posters, ZIP archives, and logs. The import pipeline is executed asynchronously by a worker process that polls for pending jobs, recursively enumerates files under each selected directory, filters supported media types, computes SHA-256 hashes for duplicate detection, and compares candidate assets to previously imported files. Duplicate files are not re-imported; instead, the software records a duplicate event linked to the corresponding import job. Non-duplicate files are copied into managed immutable storage and then processed to extract metadata and generate derived representations for rapid visual browsing.

To derive quantitative image descriptors from imported public datasets, we implemented an RGB line-profile and whole-image feature extraction workflow. For each imported image, RGB values were sampled along four canonical lines: the main diagonal, anti-diagonal, middle horizontal line, and middle vertical line. Each line was resampled to a fixed number of points and summarized by channel-wise mean, standard deviation, minimum, maximum, quartiles, median, intensity statistics, and line-profile gradient measures. In parallel, whole-image RGB, HSV, and CIE Lab histograms were computed together with global color statistics, tile-based regional mean color features on a regular grid, and simple texture descriptors including gradient mean, gradient dispersion, edge density, intensity entropy, and global contrast. Both compact summary descriptors and raw resampled line-profile vectors were exported as per-image feature matrices for downstream dimensionality reduction and clustering analyses. Overlay previews marking the sampled lines were generated for quality control and visual interpretation of the sampling geometry.

For downstream analysis, feature matrices were standardized and optionally normalized prior to dimensionality reduction; principal component analysis (PCA) was used to capture dominant variance structure, and uniform manifold approximation and projection (UMAP) was applied to visualize dataset-level organization and separation of image collections.

Image metadata are extracted with Sharp, while video metadata are obtained through ffprobe. For still images, PixelDeck generates thumbnail and preview JPEGs sized for efficient interface rendering. For videos, the worker creates a poster frame, a short preview clip, and a thumbnail based on the poster image. Each successfully processed file is inserted into the metadata store and added to the full-text search index. Export operations are handled similarly through asynchronous jobs that package selected original files into ZIP archives, preserving unique output names when needed. This background-processing model decouples long-running file operations from interactive user activity and maintains interface responsiveness during large imports and exports. A schematic overview of the PixelDeck software architecture and processing workflow is shown in **Fig. 1**.

**Figure 1.**
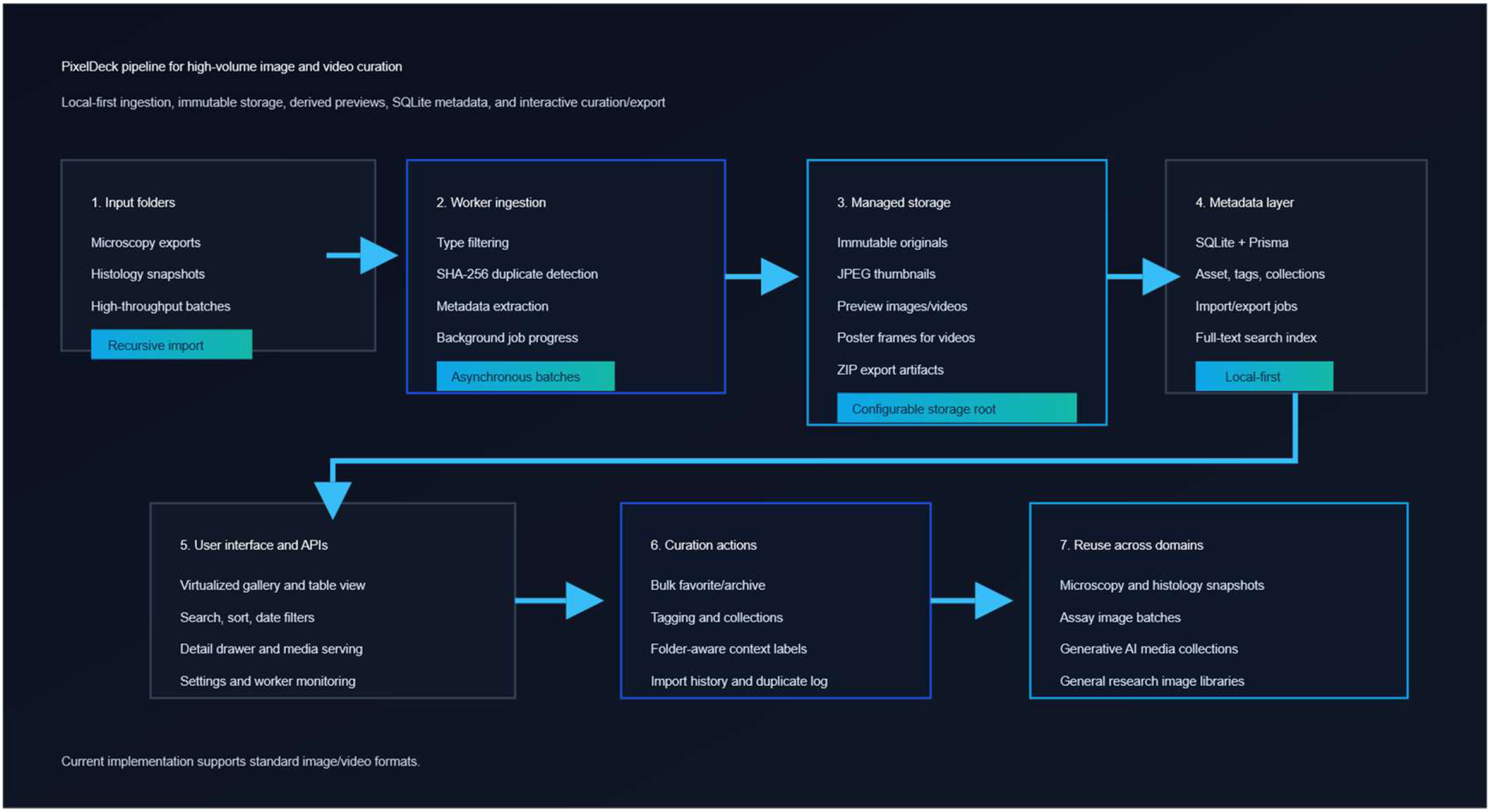
PixelDeck architecture and processing pipeline. External imaging folders are imported into managed local storage, processed by a background worker for duplicate detection, metadata extraction, derivative generation, and indexing, and then exposed through the browser interface for browsing, curation, and export.

## RESULTS

We initiated benchmark imports for public histopathology subcollections spanning PanopTILs, SICAPv2, and PanNuke datasets. These sources include RGB image collections, allowing PixelDeck to be evaluated on heterogeneous public material resembling the exported review assets commonly encountered in digital pathology and computational histology workflows. The main PixelDeck browsing interface is shown in **Fig. 2**.

**Figure 2.**
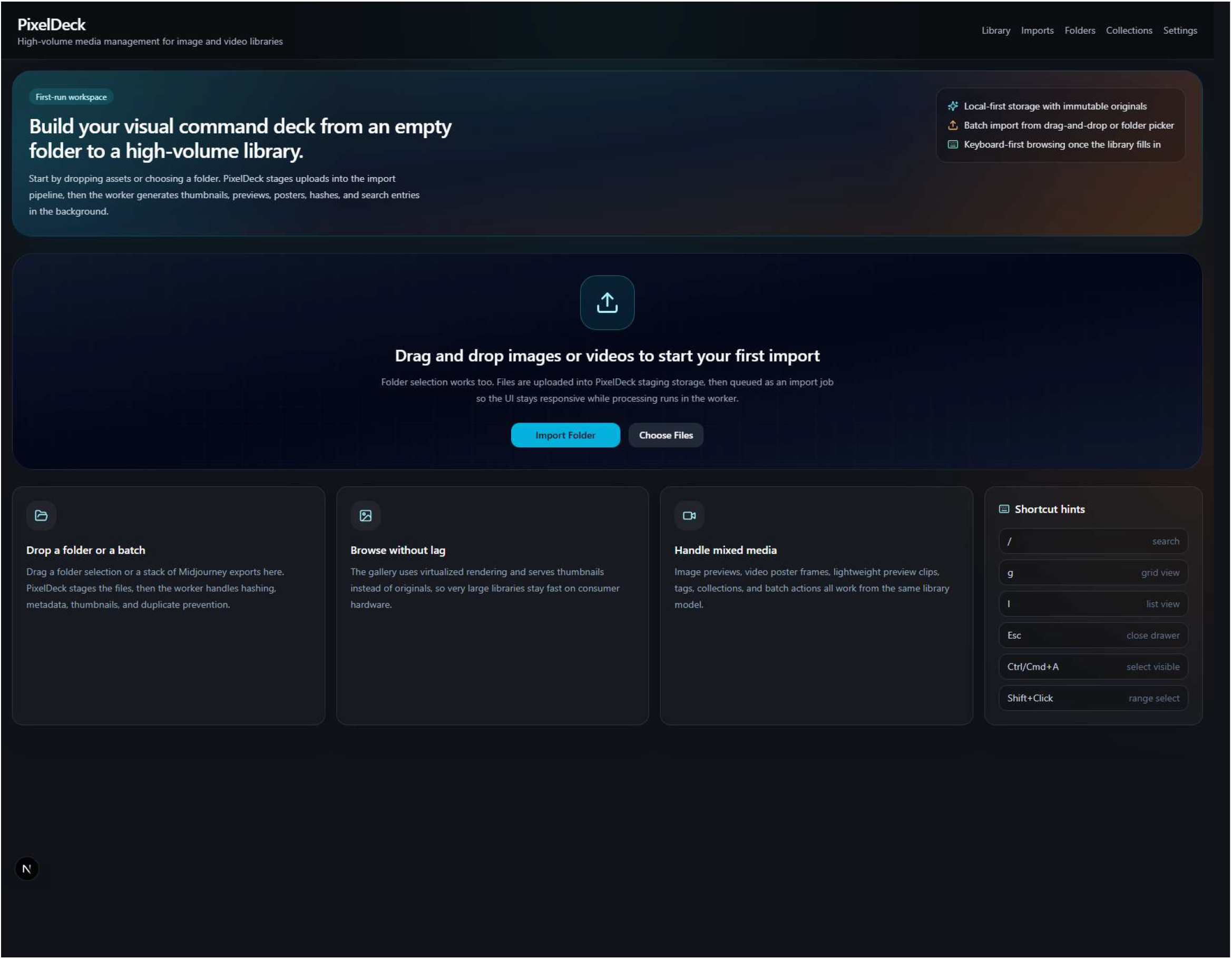
PixelDeck landing page for browsing biomedical imaging collections. The interface shows the search and filtering panel, virtualized grid layout, density controls, selection tools, and the main browsing environment used to explore imported histology, fluorescence microscopy, and high-throughput imaging assets.

Completed benchmark imports demonstrate both duplicate-aware and novel ingestion behavior across public histopathology datasets. The PanopTILs RGB subcollections completed without duplicates, skipped files, or failures, indicating clean end-to-end ingestion of previously unseen image data. In contrast, SICAPv2 and PanNuke imports showed distinct ingestion patterns consistent with their independent origins, with all files processed as novel assets.

The completed imports provide analyzable image content beyond ingestion metrics. PixelDeck-derived feature extraction shows that PanopTILs, SICAPv2, and PanNuke exhibit distinct patterns in image structure and scale. The feature pipeline transforms imported images into numerical representations, including RGB line summaries, resampled line vectors, color histograms, regional color descriptors, and texture features. These features enable dimensionality reduction using PCA and UMAP, supporting visualization and clustering of datasets by source and underlying image characteristics.

To test whether heterogeneous PanopTILs subgroups were obscuring dataset-level structure, we repeated the merged embedding analysis using only PanopTILs RGB tiles alongside SICAPv2 and PanNuke image datasets. This filtered analysis included 14,211 images in total (3,026 PanopTILs RGB, 5,000 SICAPv2, and 6,185 PanNuke) and 2,353 combined feature dimensions. Under this restricted comparison, the three datasets separated more clearly than in the earlier mixed PanopTILs runs. PCA revealed strong dataset-level structure, with PC1 explaining 75.6% of total variance, and UMAP similarly resolved three distinct clusters. These results indicate that the previously observed PanopTILs overlap was driven primarily by internal heterogeneity within the PanopTILs collection, rather than by limitations of the feature representation in distinguishing PanopTILs RGB tiles from SICAPv2 and PanNuke image sets. The filtered three-group embedding analysis is shown in **Fig. 3**.

**Figure 3.**
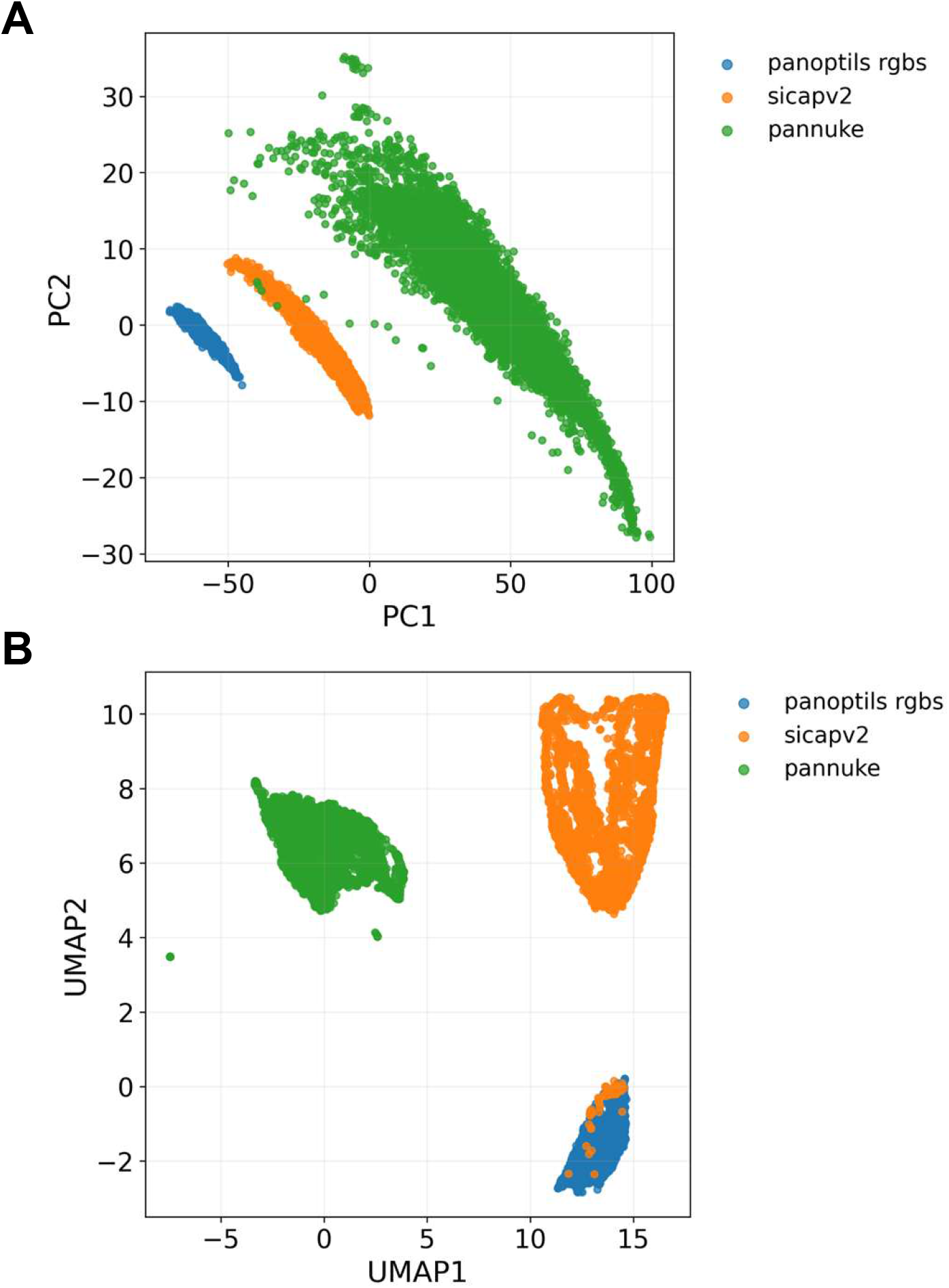
Separation of filtered RGB histopathology datasets in quantitative image-feature space. (**A**) PCA of combined image descriptors. (**B**) UMAP of the same filtered dataset using cosine distance (n_neighbors=10, min_dist=0.01). The final analysis included 3,026 PanopTILs RGB tiles, 5,000 SICAPv2 image tiles, and 6,185 PanNuke image tiles.

These analysis outputs complement the media-management functionality of PixelDeck. Import-job records quantify ingestion behavior, duplicate handling, and queue state, while feature matrices derived from imported images provide the substrate for downstream exploratory analysis and visualization. Together, they position PixelDeck not only as a local curation tool but also as a lightweight platform for quantitative characterization of heterogeneous histopathology image collections.

## DISCUSSION

PixelDeck addresses a common but under-served problem in biomedical imaging: management of large derived media collections after acquisition and analysis have already taken place. Existing platforms such as OME/OMERO, Icy, Bisque, and related systems have made foundational contributions to image data modeling, multidimensional image management, and integrated analysis environments^4, 8-11^. However, many research groups need a lighter solution for everyday local curation of exported image and video assets on workstation-class hardware. PixelDeck was designed for this specific operational space, offering an alternative to both ad hoc filesystem browsing and more infrastructure-heavy image-management systems.

The software has several practical strengths. First, its local-first design keeps both metadata and media files transparent and easy to inspect, back up, and migrate. Second, the explicit separation of immutable originals from generated browsing derivatives preserves data integrity while supporting responsive interaction. Third, asynchronous worker-based processing enables duplicate-aware import and export without blocking the user interface. Together, these design choices make PixelDeck suitable for laboratories that want a reproducible, lightweight, and open-source solution for media organization without adopting a full institutional platform.

At the same time, the current implementation has some limitations. PixelDeck is optimized for exported, review-friendly image and video formats rather than raw acquisition data, native multidimensional microscopy containers, or whole-slide pathology image formats. The software is therefore not intended to replace dedicated digital pathology platforms, image archive systems, or integrated analysis environments. In addition, the present local SQLite design is best matched to single-user or small-group deployments rather than large, concurrent, multi-user installations, although the architecture is compatible with future migration toward a more scalable backend.

## CONCLUSION

PixelDeck is an open-source, local-first media library manager for biomedical imaging workflows. By combining recursive import, content-based duplicate detection, metadata extraction, generated derivative media, full-text search, and asynchronous export in a browser-based interface, the software provides a practical way to organize and reuse large collections of exported images and videos on commodity workstations. PixelDeck is particularly relevant for histology, fluorescence microscopy, and high-throughput imaging workflows in which derived outputs must be reviewed, curated, and repurposed after primary acquisition or analysis. The current implementation offers a lightweight and transparent solution for this problem and establishes a foundation for future extensions in scientific image management.

## AVAILABILITY AND IMPLEMENTATION

Project home page: https://github.com/KidderLab/PixelDeck. Operating system(s): platform independent, with current workflow emphasis on Windows-based local deployments. Programming language: TypeScript. Other requirements: Node.js 20 or greater, pnpm 9 or greater, SQLite, FFmpeg, and ffprobe. License: MIT. Current version described in this manuscript: v0.1.0.

## ACKNOWLEDGEMENTS

This work utilized the Wayne State University High Performance Computing Grid for computational resources (https://www.grid.wayne.edu/).

## DATA AVAILABILITY

No patient data are included in this manuscript. The benchmark evaluation uses public histopathology subcollections curated from PanopTILs, SICAPv2, and PanNuke datasets. The software source code is available from the project repository.

